# Differential activation of Fyn kinase distinguishes saturated and unsaturated fats in mouse macrophages

**DOI:** 10.1101/157529

**Authors:** Elena Tarabra, Ting-Wen An Lee, Victor A Zammit, Manu Vatish, Eijiro Yamada, Jeffrey E Pessin, Claire C Bastie

**Affiliations:** Department of Medicine and Molecular Pharmacology, Albert Einstein College of Medicine, Bronx, NY, USA; Division of Biomedical Sciences, Warwick Medical School, University of Warwick, Coventry, UK; Nuffield Department of Obstetrics & Gynaecology, University of Oxford, Oxford, UK; Pediatric Endocrinology, The Valley Hospital, Ridgewood, NJ, USA; Department of Medicine and Molecular Science, Gunma University Graduate School of Medicine, Maebashi, Japan

**Keywords:** Fyn Kinase, Macrophages, Fatty acids, Adipose Tissue, Obesity

## Abstract

Diet-induced obesity is associated with increased adipose tissue activated macrophage numbers. Yet, how macrophages integrate fatty acid (FA) signals remains unclear. We previously demonstrated that Fyn deficiency (FynKO) protects against high fat diet-induced adipose tissue macrophage accumulation. Herein, we show that inflammatory markers and reactive oxygen species are not induced in FynKO bone marrow-derived macrophages exposed to the saturated FA palmitate, suggesting that Fyn regulates macrophage function in response to FA signals. Saturated palmitate activates Fyn and re-localizes Fyn into the nucleus of RAW264.7, J774 and wild-type bone marrow-derived macrophages. Similarly, Fyn activity is increased in cells of AT stromal vascular fraction of high fat-fed control mice, with Fyn protein being located in the nucleus of these cells. We demonstrate that Fyn modulates palmitate-dependent oxidative stress in macrophages. Moreover, Fyn catalytic activity is necessary for its nuclear re-localization and downstream effects, as Fyn pharmacological inhibition abolishes palmitate-induced Fyn nuclear redistribution and palmitate-dependent increase of oxidative stress markers. Importantly, mono-or polyunsaturated FAs do not activate Fyn, and fail to re-localize Fyn to the nucleus. Together these data demonstrate that macrophages integrate nutritional FA signals via a differential activation of Fyn that distinguishes, at least partly, the effects of saturated versus unsaturated fats.

## INTRODUCTION

Obesity, the most potent trigger for the development of type 2 diabetes and insulin resistance, has reached pandemic proportions and will affect more than 475 million individuals worldwide in the next 15 years [1]. Obesity is the result of dysregulation of a number of biological processes, including the disturbance of whole body homeostasis but also immune function [2]. In this regard, obesity is associated with systemic low-grade inflammation, also called meta-inflammation characterized by accumulation of immune cells in peripheral tissues such as liver, skeletal muscle and adipose tissue [3–6]. In addition to the increased immune cell number, obesity-induced inflammation is defined by a modification of immune cell populations with increased B [6] and T lymphocytes [7], neutrophils [8] and mast cells [9] and decreased natural killer cells and eosinophil numbers, promoting insulin resistance, particularly in the adipose tissue of obese individuals [10]. Interestingly, meta-inflammation shares similar sets of signaling pathways with classical inflammation. However, the hallmark of meta-inflammation is that the inflammatory processes are stimulated by the same excessive nutrient supply which also dysregulates energy metabolism and insulin sensitivity.

Remarkably, macrophages can represent up to 40% of the adipose tissue cell population in morbidly obese patients [11] and there is substantial evidence that they are involved in the establishment of inflammation-associated insulin resistance in this tissue. However, it is still under scrutiny whether the sole increase in macrophage numbers [12] or a switch in their phenotype from anti- to pro-inflammatory [13, 14] is responsible for this inflammation. Recently, presence of macrophages distinct from the M1 pro-inflammatory phenotype and resemble the M2 phenotype have been described in adipose tissues of obese humans and mice [14]. Interestingly these macrophages display a unique set of markers particularly involved in lipid metabolism that are induced by the saturated FA palmitate, demonstrating that high fat diets induce a change in macrophage phenotype and function.

Determining the molecular mechanisms that trigger these changes under a high fat diet regime is likely to offer insights into the metabolic consequences of obesity. However, how cells “sense” fatty acid (FA) cues remains to be clarified. In this regard, recent studies have focused on molecules that could provide a link between FA sensing and immune function. AMP-activated protein kinase (AMPK) has recently being suggested to mediate some of the anti-inflammatory effects of unsaturated FAs in macrophages [15, 16] by decreasing pro-inflammatory gene expression [17, 18]. Recently, the Nuclear-factor erythroid2-related factor 2 (Nrf2), a transcription factor playing physiological roles in oxidative stress has emerged as an important factor in the orchestration of the effects of FAs on macrophage function [19]. Indeed, high fat feeding inhibits Nrf2 downstream targets [20]. Interestingly, Fyn tyrosine phosphorylates Nrf2, promoting Nrf2 nuclear exclusion and subsequent degradation [21-23].

The Src family of non-receptor tyrosine kinase member Fyn is an important molecule in the integration of metabolic and inflammatory pathways. Its role as a modulator of innate immune function is well established, and we have demonstrated that Fyn is a key regulator of glucose and energy homeostasis [24]. Consistently with the role of Fyn in lipid metabolism, we have demonstrated that Fyn-deficient (FynKO) mice display higher rates of long chain FA oxidation as a result of enhanced LKB1 and AMPK activities in adipose tissue and skeletal muscle [25]. More recently, we reported that Fyn deficiency protects animals from diet-induced insulin resistance despite induction of obesity [26]. Importantly, we showed that macrophage numbers were greatly reduced in the adipose tissue of FynKO animals after high fat diets [26]. This suggested that Fyn might regulate macrophage function in response to FAs. Therefore, in this report, we investigated 1) the role of Fyn kinase in saturated FA-induced macrophage activation and 2) the effects of saturated and unsaturated FAs on Fyn function and downstream signaling.

## RESULTS

### Increased Fyn activity in adipose tissue stromal vascular fraction of high fat-fed mice

High fat diet-fed FynKO mice display reduced systemic inflammation and decreased macrophage numbers in the stromal vascular fraction (SVF) in their adipose tissue [26], suggesting that Fyn participates in the deleterious effects of high fat diets on inflammatory processes. Interestingly, Fyn protein expression was significantly increased in isolated adipose tissue macrophages (ATMs) from wild type mice following 8 weeks of high fat diet (60% Kcal) (Fig.1A, B). Consistent with this increase, Fyn tyrosine activity was increased more than 2-fold in the SVF of adipose tissue of fat-fed mice (Fig. 1C), suggesting that dietary fat content might regulate Fyn activity.

**Figure 1.**
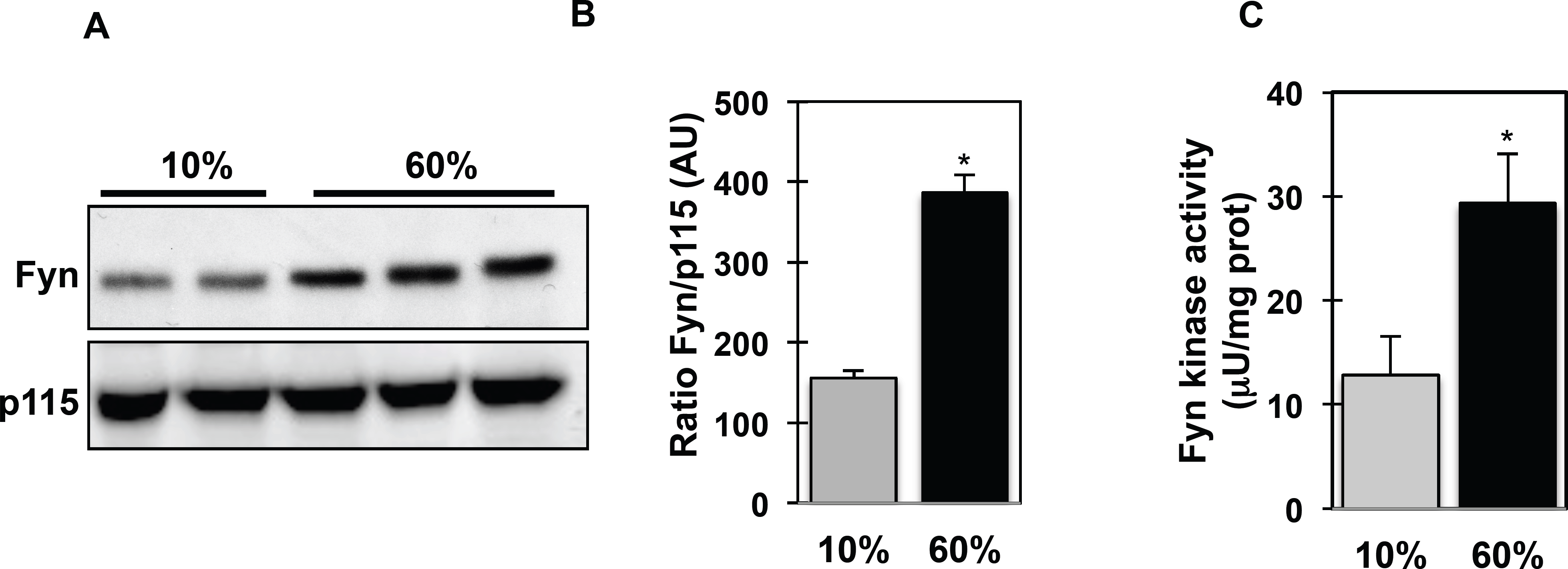

### Inflammation markers and ROS are reduced in palmitate-treated FynKO macrophages

To investigate the mechanisms by which Fyn expression/activity affects the macrophage response to FA, we first assessed inflammatory marker expression in isolated ATMs from high fat-fed control and FynKO mice. Consistent with our previous report [26], TNFα and IL6 expression were reduced in macrophages from high fat-fed FynKO mice (Fig. 2A). Importantly, whilst palmitate increased inflammatory markers (NOS2, IL6 and TNFα) in bone marrow-derived macrophages (BMDMs) from low fat-fed control mice, these markers were not induced in BMDMs of low fat-fed FynKO mice (Fig. 2B). Using shRNA-mediated knock down, we silenced Fyn in RAW264.7 macrophages (shFyn-RAW264.7) (Fig. 2C). The reactive oxygen species (ROS) production that accompanies the early stage of inflammatory processes was increased in RAW264.7 macrophages after incubation with palmitate, but not in shRNAFyn-RAW264.7 macrophages (Fig. 2D). Similarly, NAD(P)H oxidase expression, the enzyme catalyzing the production of superoxide, was not induced in Fyn-deficient macrophages after palmitate exposure (Fig. 2E). These data suggest that lack of Fyn kinase expression in macrophages is required to lessen the inflammatory effects mediated by saturated fatty acids.

**Figure 2.**
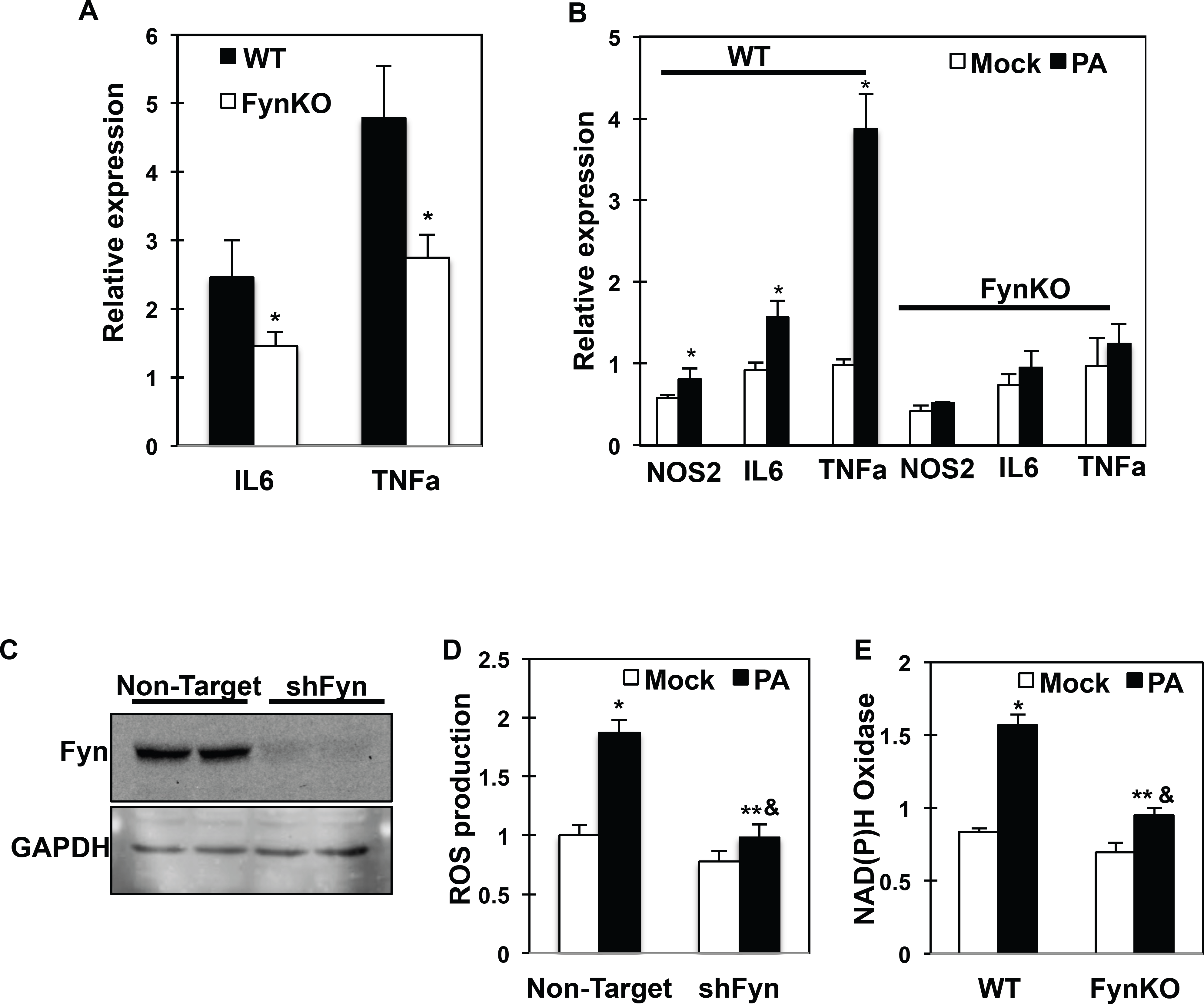

### Saturated but not unsaturated FAs redistribute Fyn in the nucleus of macrophages

Fyn subcellular localization varies from cytoplasmic to nuclear depending on which signals cells were exposed to [22, 27]. Interestingly, Fyn protein expression was increased in adipose tissue SVF of high fat-fed mice (Fig. 3A), consistent with increased Fyn activity (Fig. 1C). Also, Fyn protein expression was increased in the nuclear fraction of SVF from high fat-fed mice while Fyn signal was higher in the cytoplasm fraction of SVF from low fat-fed mice (Fig. 3A). Together, these data suggested that high fat-diet re-localized Fyn into the nucleus of cells. Consistent with this, endogenous Fyn was predominantly localized in the cytoplasm of mock-stimulated RAW264.7 cells but was re-distributed to the nucleus following palmitate exposure (Fig. 3B, panels a-d). Similar results were obtained with wild type BMDMs (Fig.3B, panels e-h) and the J774 macrophage cell line (Fig.3B, panels i-l). The percentage of cells with Fyn signal predominantly in the nucleus in each cell line is presented in Supplementary Appendix 1A, B and C.

**Figure 3.**
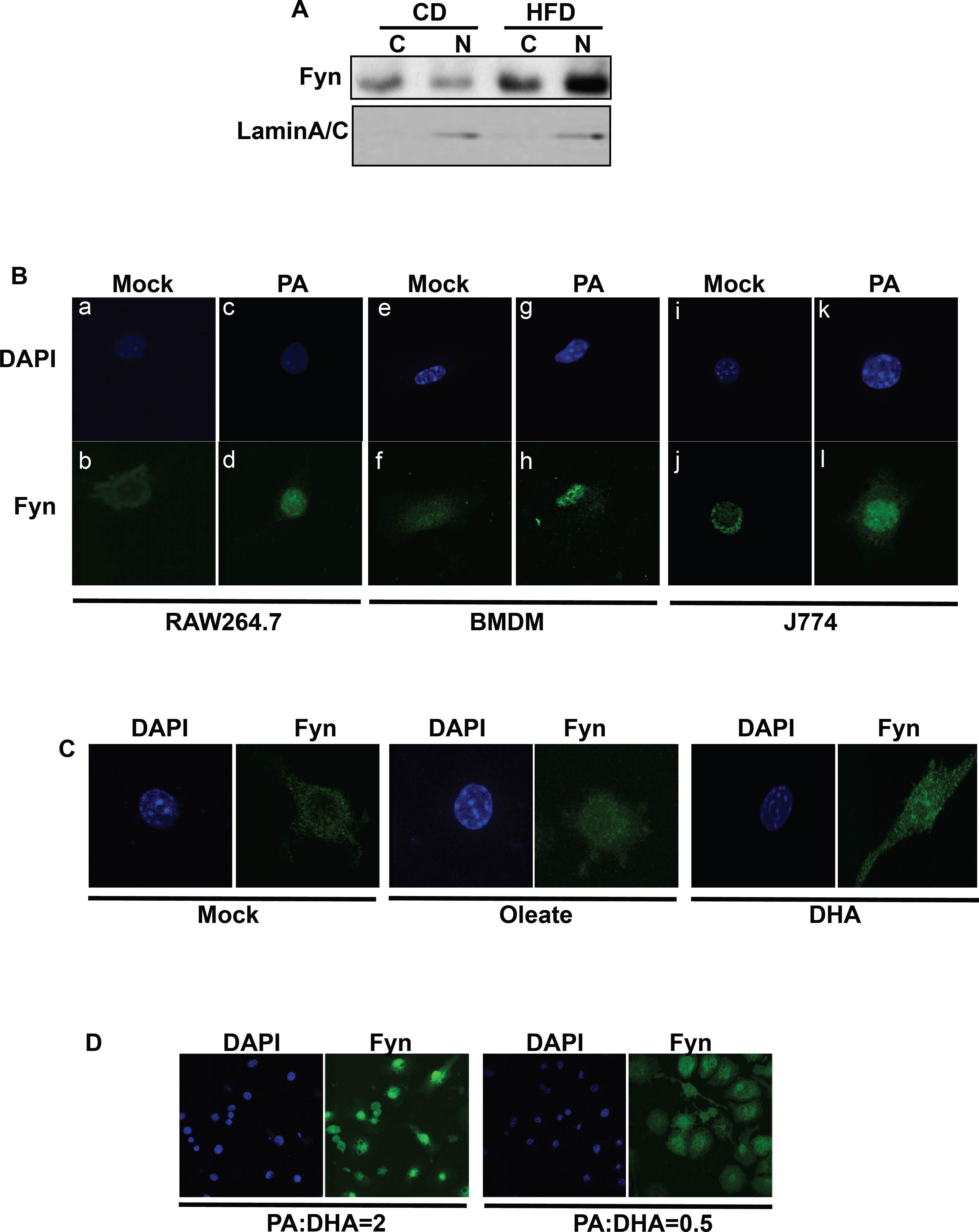

Interestingly, Fyn re-distribution to the nucleus appeared to be restricted to palmitate treatment, as Fyn remained in the cytoplasm in about 70% of cells treated with the mono-unsaturated FA oleate and in approximately 85% of cells exposed to polyunsaturated FAs docosahexaenoic acid (DHA) (Figure 3C and Supplementary Appendix 2A) or eicosapentaenoic acid (data not shown). In line with this, subcellular fractionation of RAW264.7 cells confirmed Fyn localization in nuclear fractions after palmitate exposure but cytoplasmic localization after incubation of the cells with DHA (Supplementary Appendix 2B). Interestingly, whereas in cells exposed to a mixture of saturated (palmitate) and unsaturated (DHA) FAs at a ratio similar to that found in high fat diets (PA:DHA ratio = 2), Fyn was localized to the nucleus in approximately 80% of the cells (Fig.3D), only 20% of the cells presented Fyn in the nucleus (Fig.3D) when they were incubated in a mixture where DHA concentration was higher than this of palmitate (PA:DHA ratio = 0.5). These results are essentially consistent with the fact that Fyn protein can be found in the nucleus of SVF from low-fat fed mice (Fig.3A), as the mice are also exposed, although in lesser amount, to saturated fats in their diet.

### Saturated but not unsaturated FAs activate Fyn kinase in macrophages

Importantly, palmitate-induced Fyn nuclear re-localization was abolished when cells were treated with the Fyn kinase inhibitor SU6656 (Fig.4A, panels g, h and Fig. 4B) suggesting that Fyn kinase catalytic activity is required for the observed palmitate-mediated increase in Fyn nuclear localization. Supporting this, Fyn kinase activity was 6-fold higher in cells treated with palmitate, whilst it was much more moderately increased, if at all, after oleate (2-fold) or DHA (no increase) exposure (Fig. 4C).

**Figure 4.**
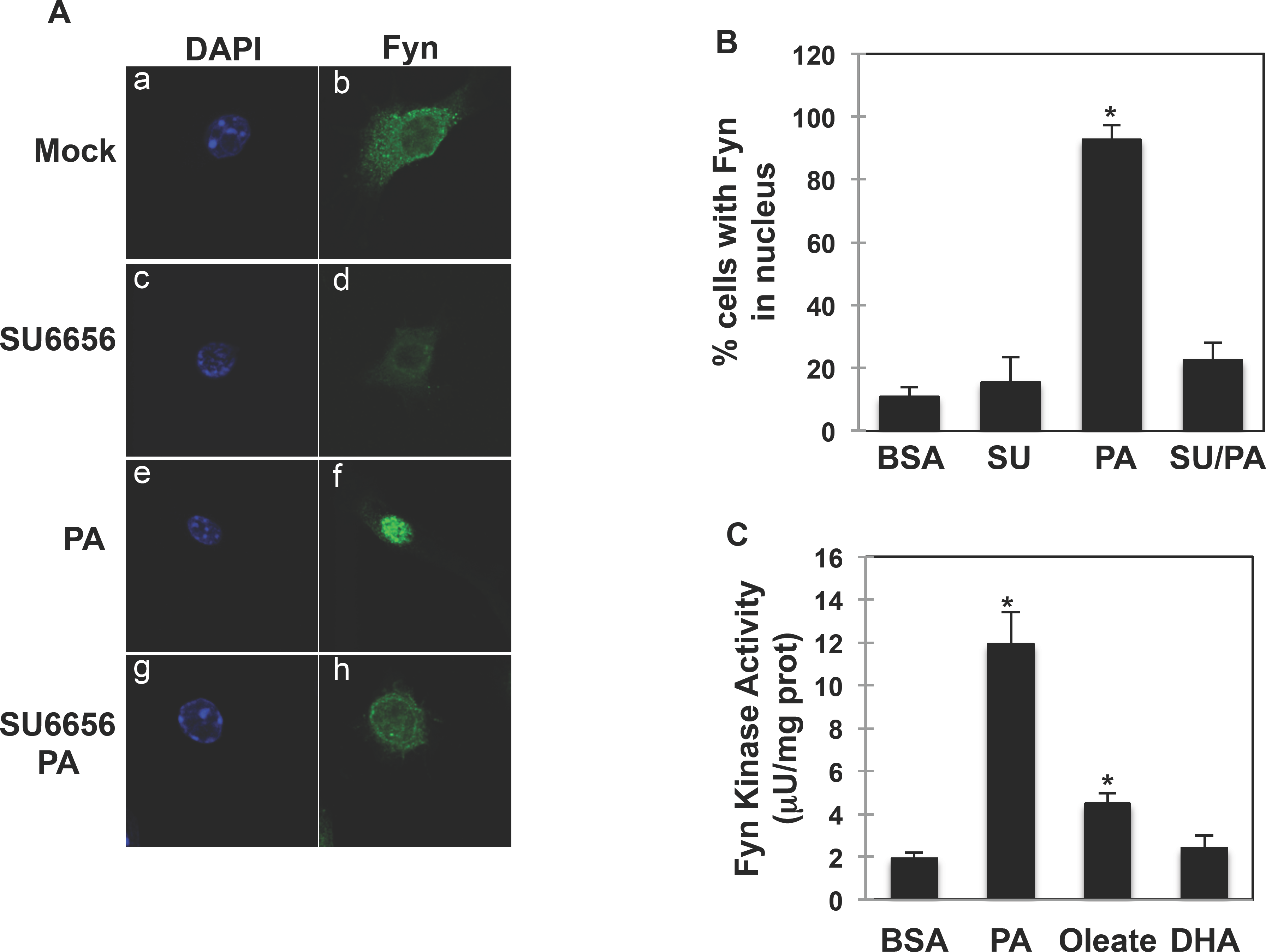

### Fyn inhibition blocks palmitate pro-inflammatory effects

To investigate the physiological relevance of Fyn subcellular re-localization after palmitate exposure, and whether this affects macrophage activation in response to saturated fats, we next investigated palmitate effects on Nrf2 subcellular localization. Nrf2 is an important transcription factor that regulates anti-oxidative gene expression and is under Fyn regulatory control. Nrf2 localized in the cytoplasm of unstimulated RAW264.7 macrophages (Fig. 5A) as previously reported [28-30]. Time course experiments in RAW264.7 macrophages treated with palmitate revealed that both Fyn and Nrf2 accumulated in the nucleus after a short palmitate exposure (Fig.5A). However, while Fyn remained in the nucleus (Fig. 5A, middle panels), the Nrf2 signal decreased in the nucleus with time (Fig. 5A, right panels) and Nrf2 was cytosolic in approximately 75 % of cells after 10 minutes (Fig. 5A and Supplementary Appendix 3). This was essentially confirmed by subcellular fractionations of RAW264.7 cells (Fig.5B). Notably, Nrf2 expression appeared to decrease after 60 min (Fig. 6B, lower panel), which could be explained by increased Keap1 expression and correlated increase in Nrf2 ubiquitination levels observed after 30 min (Supplementary Appendix 4.). To evaluate Nrf2 functionality after palmitate exposure, Nrf2 downstream targets NAD(P)H dehydrogenase [quinone]-1 (Nqo-1) and Heme oxygenase-1 (HO-1) mRNA and protein expression were determined in RAW264.7 cells. Consistently with Nrf2 cytosolic localization, Nqo-1 mRNA expression was reduced after palmitate treatment and importantly Nqo-1 expression was rescued after Fyn inhibition with SU6656 (Fig. 5C). Similarly, HO-1 protein expression was reduced after palmitate treatment and rescued after Fyn inhibition with SU6656 (Fig.5D, E) in RAW264.7 as well as in J774 (Supplementary Appendix 5) macrophage models.

**Figure 5.**
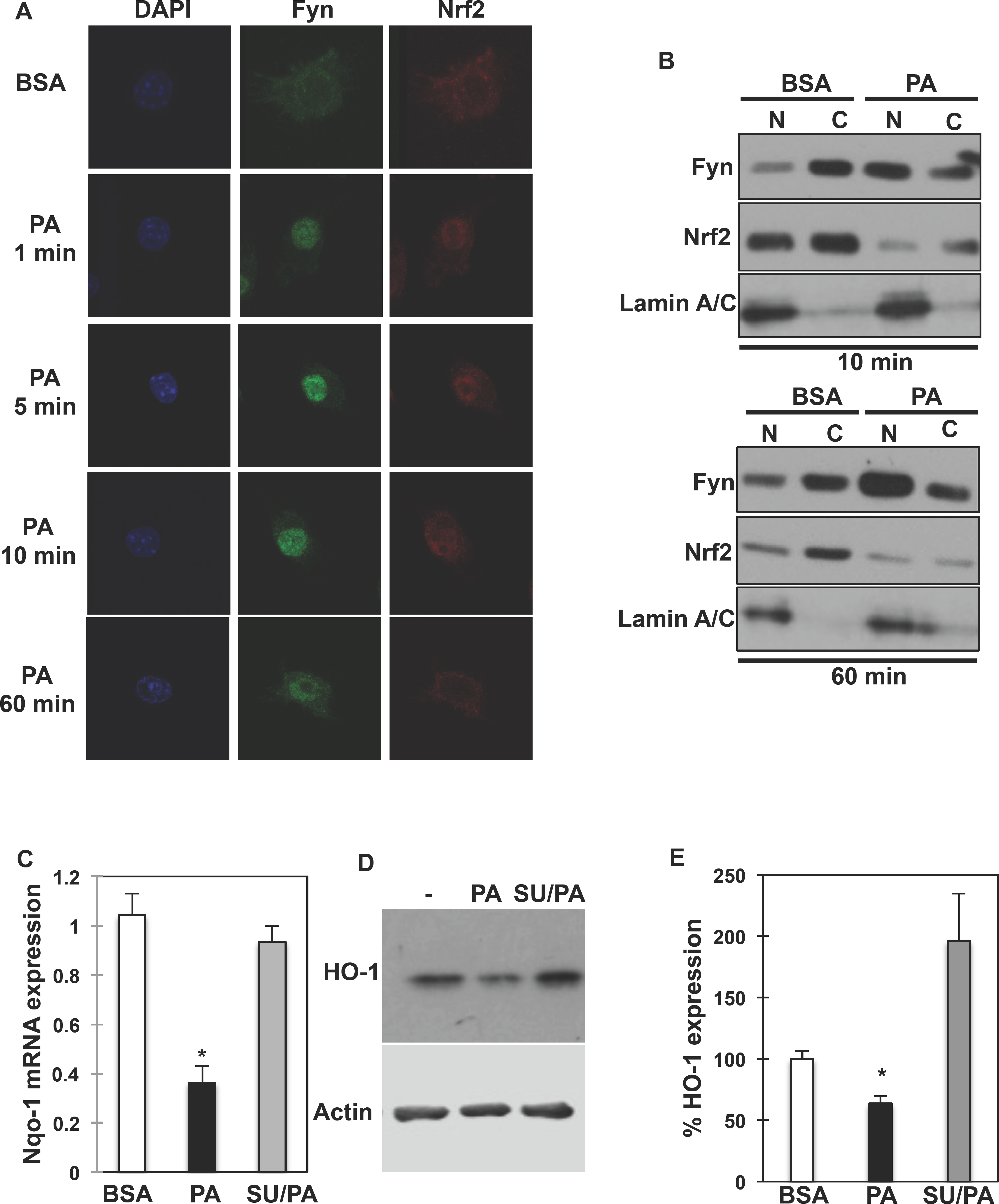

## DISCUSSION

Obesity is a major worldwide health dilemma aggravated by refined diets rich in fats. Obesity is characterized by chronic low-grade inflammation and high fat diets promote invasion of immune cells, particularly macrophages into peripheral tissues. Recent studies have suggested that the number of macrophages [12] rather than their phenotype increases inflammation in obese individuals. This is also supported by the recent observation that M1 cell-surface markers are lacking in ATMs of obese humans and mice [14], which demonstrates that the phenotypic switch from anti-inflammatory (M2) to pro-inflammatory (M1) macrophages, and its role in adipose tissue inflammation is more complex than originally described. However, Kratz et al. [14] also showed that the saturated FA palmitate stimulates the macrophage population to resemble the M2 phenotype, e.g. by having an increased lipogenic profile. This strongly supports the concept that high-fat diets participate in the modification of macrophages activation in peripheral tissues. However, not all fats are equal. Indeed, saturated fatty acids (FAs) increase pro-inflammatory gene expression, cytokine expression and reactive oxygen species (ROS) production in macrophages [31, 32] while unsaturated, particularly n-3 poly-unsaturated fatty acids (PUFAs) reduce pro-inflammatory processes in macrophages [33]. While the effects of FAs on metabolic cascades (e.g. insulin signaling) have been largely studied, the role of these nutrients and how they modulate the immune function is still unclear. Particularly, the molecular links (or “sensors”) relaying FA effects on macrophages function are still unknown. Therefore, molecules linking immune function and energy are of special interest. Previously, we showed that Fyn is a regulator of whole body lipid metabolism [24]. Remarkably, Fyn is also a major regulator of innate immune function, participating in the Toll-Like Receptor (TLR) signaling in macrophages as well as the T-cell receptor in T cells [34]. Additionally, Fyn interacts with the FA transporter CD36 [35, 36], which expression is greatly induced in palmitate-treated macrophage [14], therefore we hypothesized that Fyn could transduce the inflammatory effects of saturated FAs. Consistently with this, we determined that Fyn kinase activity was greatly increased in response to palmitate and palmitate pro-inflammatory effects were blunted in BMDMs lacking Fyn. Interestingly, the polyunsaturated FA (DHA) was unable to activate Fyn and Fyn was not re-localized to the nucleus in response to DHA (Fig. 3 and Fig. 4), which suggests that Fyn activation might be part of a signaling cascade distinguishing between the effects of saturated and unsaturated FAs on macrophage function. Further studies are necessary to identify mechanisms by which saturated and unsaturated FAs have divergent effects of Fyn kinase activation. It is suggested that Fyn distributes diffusely throughout the plasma membrane in some macrophage models and under particular stimuli, Fyn may undergo endocytosis [37]. As such, one could argue that saturated but not unsaturated FAs might trigger such mechanism to induce Fyn re-localization. It is known that CD36 and Fyn kinase interact [35], thus it is possible that mechanisms leading to palmitate-dependent Fyn activation might be mediated by CD36, which is induced by palmitate in macrophages. Nonetheless the molecular mechanism of activation, whether it would be an association/dissociation or an indirect mechanism, remains to be investigated and this was beyond the scope of the current study.

We found that, following palmitate exposure, Fyn rapidly re-localizes into the nucleus of macrophages. This was subsequently followed by Nrf2 nuclear exclusion. Interestingly, Nrf2 action is also regulated by high fat diets, and FA-dependent regulation of Nrf2 appears to be dependent on the nature of dietary fats [38], i.e. saturated fats inhibit Nrf2 signalling whereas unsaturated fats activate this pathway [20]. Therefore, it appears that FAs regulate Nrf2 function in an opposite pattern to that through which they regulate Fyn activity, as Fyn exerts an inhibitory action on Nrf2. Our data and those of others show that, under basal conditions, Nrf2 localizes in the cytoplasm of cells [28-30]. However, Nrf2 quickly re-localizes to the nucleus after palmitate exposure, along with Fyn kinase. This might initially appear counter-intuitive since Fyn inhibits Nrf2 action by promoting Nrf2 nuclear exclusion. However, we observed that after this first phase, the Nrf2 nuclear signal gradually decreased whereas Fyn remained in the nucleus, suggesting that Nrf2 function was inhibited as a consequence of the nuclear localization of Fyn. In line with this, we observed a decrease in the expression of downstream targets of Nrf2 after palmitate exposure of the cells. Importantly, the effects of palmitate were blocked when Fyn was inhibited by SU6656, suggesting that Fyn kinase activity is necessary for palmitate-dependent inhibition of Nrf2 pathway.

Mechanisms leading to Nrf2 initial nuclear localization in response to palmitate remain to be elucidated, however, it appears reasonable to suggest that this could be a mechanism to protect the cells from the deleterious effects on ROS production. Supporting this, we observed that expression of the Nrf2 targets Nqo-1 and HO-1 were transiently elevated shortly after palmitate exposure (data not shown). This early phase was then inhibited by the persistent presence of Fyn in the nucleus, which triggers Nrf2 nuclear exclusion and cytosolic degradation.

Dietary fats are well-established modulators of macrophage function, however, molecular events linking FAs to macrophage activation are not completely identified. Building on our present data, we provide a novel mechanism by which macrophages respond differentially to saturated than to unsaturated

FAs through the selective activation of Fyn kinase by saturated FA, and its antagonism by unsaturated FAs. This establishes Fyn as an integral component of a signaling pathway able to sense dietary cues contributing to the establishment of inflammation in states of nutritional obesity. Further studies are now necessary to identify the upstream regulatory mechanisms by which saturated and unsaturated FAs differentially regulate Fyn kinase activity and how Fyn is successively transported to the nucleus of macrophages.

## MATERIAL AND METHODS

### Reagents

Rabbit polyclonal Fyn (FYN3), Heme Oxygenase-1, Nrf2 (H300) and goat polyclonal Nrf2 (T19) were from Santa Cruz Biotechnology (Santa Cruz, CA, USA) and GAPDH antibody was from MBL international (Woburn, MA,USA). LaminA/C and P115 antibodies were from Cell Signaling Technology (Danvers, MA, USA).

### Animals

pp59^*fyn*^ knockout mice (FynKO) (129-Fyn ^tm1/sor^ /J) and their controls <u>(002448 129S1/SvImJ)</u> were obtained from the Jackson Laboratory (Bar Harbor, ME, USA) and housed in a facility equipped with a 12 h light/dark cycle. Animals were fed either a standard chow diet (Research Diets, New Brunswick, NJ, USA) containing 70% (Kcal) carbohydrates, 20 % protein, and 10 % fat or a high fat diet containing 20% (Kcal) carbohydrates, 20% protein and 60% fat for 10 weeks. Composition of the diets D12392 (60% fat) and D12450B (control diet) can be found at http://www.researchdiets.com/opensource-diets/stock-diets. Essentially, saturated fat was increased by 10% and polyunsaturated fats were reduced by 15% in D12492 compared to the control diet. All studies were approved by and performed in compliance with the guidelines of the Yeshiva University Institutional Animal Care and Use Committee (IACUC).

### Bone Marrow Derived Macrophages (BMDM) isolation and differentiation

BMDMs from control and FynKO mice were prepared using a standard method. Briefly, marrow was flushed from the femurs using DMEM media with 10% FBS. The suspension was filtered through a 70 μm cell strainer and red cells were removed. The remaining precursor cells were plated and differentiated into BMDMs with macrophage colony-stimulating factor (M-CSF, 10ng/ml) (R&D Systems, Minneapolis, MN, USA). Differentiation was typically achieved within 6 days.

### Stromal Vascular Fraction (SVF) cells isolation

Minced adipose tissues samples were treated with 0.05 mg/mL Liberase (TM Research Grade; Roche Applied Science, Indianapolis, IN) and incubated at 37°C for 20 min. Samples were passed through a sterile 250 mm nylon mesh. The suspension was centrifuged at 1,000xg for 5 min. The precipitated cells (SVF) were re-suspended in erythrocyte lysis buffer. The erythrocyte-depleted SVF cells were centrifuged at 500xg for 5 min and re-suspended in DMEM medium supplemented with 10%FCS and 1% Penicillin/Streptomycin.

### Macrophages isolation

SVF from lean and fat-fed mice (n=5 for the fat-fed mice and n=8 for the control mice) were prepared as described above. SVF were treated with erythrocyte lysis buffer and washed two times in PBS. Positive selection to enrich for F4/80 cells was performed using the MagniSort Mouse F4/80 positive selection kit (eBioscience,Inc,). Briefly, 10^8^ cells were suspended in a separation buffer and incubated with the selection antibody for 60 min. Cells were washed and selection beads were added to the suspension. Following sorting, cells were washed and protein homogenates were immediately prepared.

### Fatty acid preparation and incubation

Palmitic acid (PA) (Sigma, St. Louis, MO, USA) 50 mM was prepared in methanol and stored at −20°C. On the day of use, indicated PA concentrations were prepared in fatty acid-free and low endotoxin BSA at a ratio FA: BSA= 2. Equal volumes of methanol (final concentration < 0.1%) in fatty acid-free BSA were applied to control cells. Where Fyn pharmacological inhibitor was used, cells were treated with 5 μM SU6656 (Calbiochem, San Diego, CA, USA) for 60 min.

### Fyn tyrosine activity

Activity was determined with a Universal Tyrosine Kinase activity assay kit (Takara Bio Inc., Shiga, Japan). Cells were homogenized in a NP40-based buffer. Homogenates were centrifuged for 15 min at 14,000 xg at 4°C and supernatants were collected. Whole cell lysates (0.5-1 mg) were incubated with 4 μg of FYN3 rabbit polyclonal antibody coupled with Catch and Release columns (Millipore, Billerica, MA, USA) for 2 h at 4°C. Immuno-complexes were diluted and samples were mixed with an ATP solution before being incubated in an ELISA plate. Kinase activity was corrected by the protein concentration.

### Quantitative PCR analysis

Cells were homogenized into QIAzol Lysis Reagent. Total RNA was isolated using RNeasy^®^ Mini Kit (Qiagen Sciences, Maryland, USA) and reverse-transcribed to cDNA using the SuperScript VILO cDNA synthesis kit (Invitrogen, Carlsbad, CA, USA). TaqMan (Applied Biosystems, Branchburg, NJ, USA) RT-PCR was performed for measurement of mRNA of Tumor Necrosis Factor-alpha (TNF-a) (Mm00443260_g1), Interleukin-6 (IL-6) (Mm00446190_m1), Nitric oxide synthase-2 (Nos2) (Mm00440502_m1), Fyn (Mm00433373_m1), Heme Oxygenase-1 (HO-1) (Mm00516005_m1), NAD(P)H dehydrogenase [quinone]-1 (Nqo-1) (Mm01253561_m1),and NAD(P)H Oxidase. Relative expression levels of the mRNAs were determined using standard curves using the Applied Biosystems 7900HT Sequence Detection System from Applied Biosystems. Samples were adjusted for total mRNA content by comparison with Rpl7 expression. All primer-probe mixtures were from Applied Biosystems (Branchburg, NJ, USA).

### Western blotting

Cells were homogenized in a NP-40 lysis buffer containing protease and phosphatase inhibitors and protein concentration was determined using the BCA method. Homogenates were centrifuged for 15 min at 13,000 xg at 4°C and supernatants were collected. Protein samples (40 μg) were separated onto 8 or 10% reducing polyacrylamide gels and electroblotted onto Immobilon-P polyvinylidene difluoride membranes (Bio-Rad Laboratories, Hercules, CA, USA). Immunoblots were blocked either 5% milk in Tris-buffered saline or with Blocking Buffer for Fluorescent Western Blotting (Rockland Antibodies & Assays, Gilbertsville, PA, USA) for 2 h at room temperature and incubated overnight at 4°C with the indicated antibodies in Tris-buffered saline and 0.05% Tween 20 (TBST) containing 1% BSA. Membranes were washed in TBST and incubated with horseradish peroxidase-conjugated secondary antibodies (1:30,000) for 45 min at room temperature. Membranes were washed in TBST, and antigen-antibody complexes were visualized by chemiluminescence using an ECL kit (Pierce, Rockford, IL, USA). Alternatively, immunoblots were incubated with IRDye800CW Goat Anti Mouse (H+L) or IRDye680 Goat Anti Rabbit (H+L) secondary antibodies and signal was detected with the Odyssey^®^ Infrared Imaging System (Li-COR Biotechnology, Lincoln, NE, USA). The relative band intensity was quantified using the ImageJ software.

### Nuclear and cytoplasmic protein fractions

Cells were treated as indicated and nuclear and cytoplasm extracts were prepared using a kit from Pierce (Rockford, IL, USA). Proteins were separated onto 10% reducing polyacrylamide gels and electroblotted onto Immobilon-P polyvinylidene difluoride membranes. Immunoblots were processed as described above. Antigen-antibody complexes were visualized by chemiluminescence or using the Odyssey^®^ Infrared Imaging System, as above.

### Cell culture and silencing

The human embryonic kidney cells (HEK-293T), the mouse macrophage cell lines RAW264.7 (ATCC TIB-71) and J774 (ATCC TIB-67) were obtained by ATCC (Manassas, VA,USA) and maintained in DMEM supplemented with 10% fetal calf serum and 1% penicillin/streptomycin at 37°C in a humidified incubator with 5%CO2. Fyn kinase knock-down and control stable cell lines were produced transducing macrophage cell lines with either MISSION shRNA lentivirus Fyn particles (TRCN000023380) or MISSION non-mammalian target shRNA control particles (SHC002)(Sigma). The transduced cells were then selected with 3 μg/ml Puromycin for 10 days. Transient transfections of HEK293 cells were performed using the FugeneHD (Roche Applied Science, Basel, Switzerland) according to manufacturer’s protocol.

### Immunofluorescence

Cells were seeded in 8 wells-chamber slides and stimulated with PA, oleate, DHA or SU6656. After the indicated incubation time, cells were washed with PBS and fixed for 30 min in PBS containing 4% paraformaldehyde and 2% sucrose, followed by three washes in PBS. Slides were incubated in PBS with Triton-X-100 (PBS-T) for 5 min at 4°C to permeabilize the cells. Cells were washed and blocked in PBS-T with BSA 5% for 1 h at room temperature and incubated with either Fyn or Nrf2 antibody diluted 1/200 in PBS-T-BSA 2%, followed by Alexa Fluor 488 anti-rabbit IgG or IgG or Alexa Fluor 594 anti-goat IgG for 1 h at room temperature. Samples were mounted with Prolong Gold anti-fade reagent with DAPI (Life Technologies). Fluorescence images were acquired using a confocal fluorescence microscope (TCS SP5 confocal; Leica microsystems, Buffalo Grove, IL, USA).

### Cellular Reactive Oxygen Species Detection Assay

Cells were seeded onto 96 wells plate and pre-loaded with 10 μM 2′,7′-dichlorodihydrofluorescein diacetate (H2DCFDA) (Life Technologies) at 37°C for 60 min at room temperature. After 2 washes with PBS, reactive oxygen species (ROS) production was determinate by quantifying the fluorescence of the cells after exposure to PA for 10 min using the BioTek Synergy 2 plate reader (BioTek Instruments, Winooski, VT, USA).

### Statistics

Results are expressed as mean ± standard error of the mean (s.e.m). The data were analyzed by one-way ANOVA followed by post hoc analysis for comparisons between individual groups when group numbers were >2. Differences were considered statistically significant at a level of p<0.05. Differences between 2 treatments were tested for statistical significance (p<0.05) using a Student’s unpaired t-test.

### ABBREVIATIONS

KO: Knockout
WT: Wild Type
BMDM: Bone Marrow Derived Macrophages
HFD: high fat diet
Nrf2: Nuclear-factor erythroid2-related factor2
AMPK: AMP-activated protein kinase
DHA: docosahexaenoic acid
PA: palmitate
SVF: stromal vascular fraction
ROS: reactive oxygen species

## AUTHOR CONTRIBUTIONS

Original idea, experimental conception, manuscript writing and editing (CCB); Experimental execution, writing and editing (ET; MV, CCB); Experimental execution (TWAL, EY); writing and editing (VAZ, JEP)

## ACKNOWLEDGEMENT

The authors wish to thank the members of the Diabetes Center at Albert Einstein College of Medicine for their continuous support.

